# Immunological profiling of COVID-19 patients with pulmonary sequelae

**DOI:** 10.1101/2021.06.03.447023

**Authors:** Jianghua Wu, Lu Tang, Yanling Ma, Yu Li, Dongmei Zhang, Qian Li, Heng Mei, Yu Hu

## Abstract

Cellular immunity may be involved in organ damage and rehabilitation in patients with coronavirus disease 2019 (COVID-19). We aimed to delineate immunological features of COVID-19 patients with pulmonary sequelae (PS) one year after discharge. 50 COVID-19 survivors were recruited and classified according to radiological characteristics: 24 patients with PS and 26 patients without PS. Phenotypic and functional characteristics of immune cells were evaluated by multiparametric flow cytometry. Patients with PS had an increased proportion of natural killer (NK) cells and lower percentage of B cells compared to patients without PS. Phenotypic and functional features of T cells in patients with PS were predominated by the accumulation of CD4+ T cells secreting IL-17A, short-lived effector-like CD8+ T cells (CD27-CD62L-) and senescent T cells with excessive secretion of granzyme-B/perforin/IFN-γ. NK cells were characterized by the excessive secretion of granzyme-B and perforin and the downregulation of NKP30 and NKP46; highly activated NKT and γδ T cells exhibited NKP30 and TIM-3 upregulation and NKB1 downregulation in patients with PS. However, immunosuppressive cells were comparable between the two groups. The interrelation of immune cells in COVID-19 was intrinsically identified, whereby T cells secreting IL-2, IL-4 and IL-17A were enriched among CD28+ and CD57-cells and cells secreting perforin/granzyme-B/IFN-γ/TNF-α expressed markers of terminal differentiation. CD57+NK cells, CD4+perforin+ T cells and CD8+CD27+CD62L+ T cells were identified as the independent predictors for residual lesions. Overall, our findings unveil the profound imbalance of immune landscape that may correlate with organ damage and rehabilitation in COVID-19.

**IMPORTANCE:** A considerable proportion of COVID-19 survivors have residual lung lesions, such as ground glass opacity and fiber streak shadow. To determine the relationship between host immunity and residual lung lesions, we performed an extensive analysis of immune responses in convalescent patients with COVID-19 one year after discharge. We found significant differences in immunological characteristics between patients with pulmonary sequelae and patients without pulmonary sequelae one year after discharge. Our study highlights the profound imbalance of immune landscape in the COVID-19 patients with pulmonary sequelae, characterized by the robust activation of cytotoxic T cells, NK cells and γδ T cells as well as the deficiencies of immunosuppressive cells. Importantly, CD57+NK cells, CD4+perforin+ T cells and CD8+CD27+CD62L+ T cells were identified as the independent predictors for residual lesions.

## INTRODUCTION

As of early May 2021, more than 150 million people have developed coronavirus disease (COVID-19), a pandemic that has killed approximately 3 million people. Caused by acute respiratory syndrome coronavirus 2 (SARS-CoV-2), COVID-19 exhibited a highly variable clinical course, ranging from a high proportion of asymptomatic and mild infections to severe and fatal disease(1). With the help of interventions and immediate medical support, most patients have recovered. Nevertheless, a considerable proportion of survivors have unresolved health issues, such as pulmonary fibrosis and gas diffusion impairment(2). Thus, understanding the clinical and immunological features of convalescent individuals is critically important to elucidate the immunopathogenesis of COVID-19 and facilitate the development of effective immune interventions.

Early infection with SARS-CoV-2 can induce efficient innate immunity, followed by an adaptive immune response to control the virus(3). Such synchronized interaction between innate and adaptive immunity exquisitely mediates both viral control and host toxicity in COVID-19. Although activated immune cells orchestrate a protective function against SARS-CoV-2, they also participate in tissue damage if overactivated by inflammatory stimuli(4). Numerous studies have investigated humoral and cellular immune responses in patients who have recovered from COVID-19(5-7). For example, peripheral blood SARS-CoV-2-specific T cells are detectable in convalescent patients, especially SARS-CoV-2-specific CD8+ and CD4+ T cells, which correlated not only with serum neutralizing activities but also with disease severity(7-11). One study detected both virus-specific memory B and T cells in individuals who have recovered from mildly symptomatic COVID-19, showing that these cells not only persist, but in some cases increased numerically over three months after symptom onset(5). However, it remains unclear how cellular immunity mediates long-term protective and pathogenic inflammation in COVID-19.

Similar to influenza virus infection(12), SARS-CoV-2-mediated lung damage entails the interplay among aberrantly activated monocytes/macrophages producing IL-1β, inflammation-induced impairment of alveolar epithelial regeneration, and expansion of CTHRC1+ pathological fibroblasts that promoted fibrosis and may impair regeneration(13). Zhao et al. identified clonally expanded tissue-resident Th17 cells in the lungs of patients even after SARS-CoV-2 clearance, which may interact with profibrotic macrophages and cytotoxic CD8+ T cells leading to the formation of pulmonary fibrosis(4). Although lung lesions in COVID-19 patients could be completely absorbed during follow-up with no sequelae, residual ground-glass opacification, interstitial thickening or fibrotic-like changes were observed in most patients who survived severe COVID-19(2, 14, 15). Immunologic determinants underlying pulmonary sequelae (PS) are not fully understood in COVID-19. To determine the immunopathogenesis of PS, we recruited 24 COVID-19 survivors with PS and 26 patients without PS and performed comprehensive assessments of immunological profiling. Our study highlights the profound imbalance of immune landscape in COVID-19 patients with PS, as characterized by robust activation of cytotoxic T cells, NK cells and γδ T cells as well as deficiencies in immunosuppressive cells.

## RESULTS

### Clinical characteristics of convalescent patients with COVID-19

A total of 50 convalescent patients with COVID-19 were included and categorized as 24 patients with PS (PPSs) and 26 patients without PS (NPSs) based on radiological characteristics. As shown in Table 1, the mean age of all patients was 53.96 years, with a significantly older age for PPSs than NPSs (59.63 vs 48.73, p<0.0003). The most common comorbidities among the patients were 20.00% with hypertension, 14.00% with diabetes mellitus and 10.00% with cardiopathy. Nasopharyngeal swab SARS-CoV-2 RNA detection was negative in all patients one year after discharge (Table 1). SARS-CoV-2 IgM (2 [8.33%] patients) and IgG levels (22 [91.67%] patients) in PPSs were positive, while SARS-CoV-2 IgM (2 [7.69%] patients) and IgG levels (23 [88.46%] patients) in NPSs were positive (Table 1). The predominant patterns of PS were ground glass opacity (24 [100.00%]), fiber streak shadow (21 [87.50%]) and tractive bronchiectasis (8 [33.33%]) in PPSs (Table 1). Representative chest CT scans longitudinally exhibited the change of lung lesions (Fig. 1A). Although the lung lesions gradually resolved in all patients, ground glass opacity (GGO), fiber streak shadow, tractive bronchiectasis, reticulation and bronchovascular bundle distortion could be observed in the representative patients (Fig. 1A-C). Importantly, there was a statistically significant increment in hemoglobin in PPSs in comparison to NPSs (Table 1), suggesting that residual pulmonary lesions may influence diffusion capacity and induce a compensatory increase in hemoglobin in those with PS.

**Table 1.**
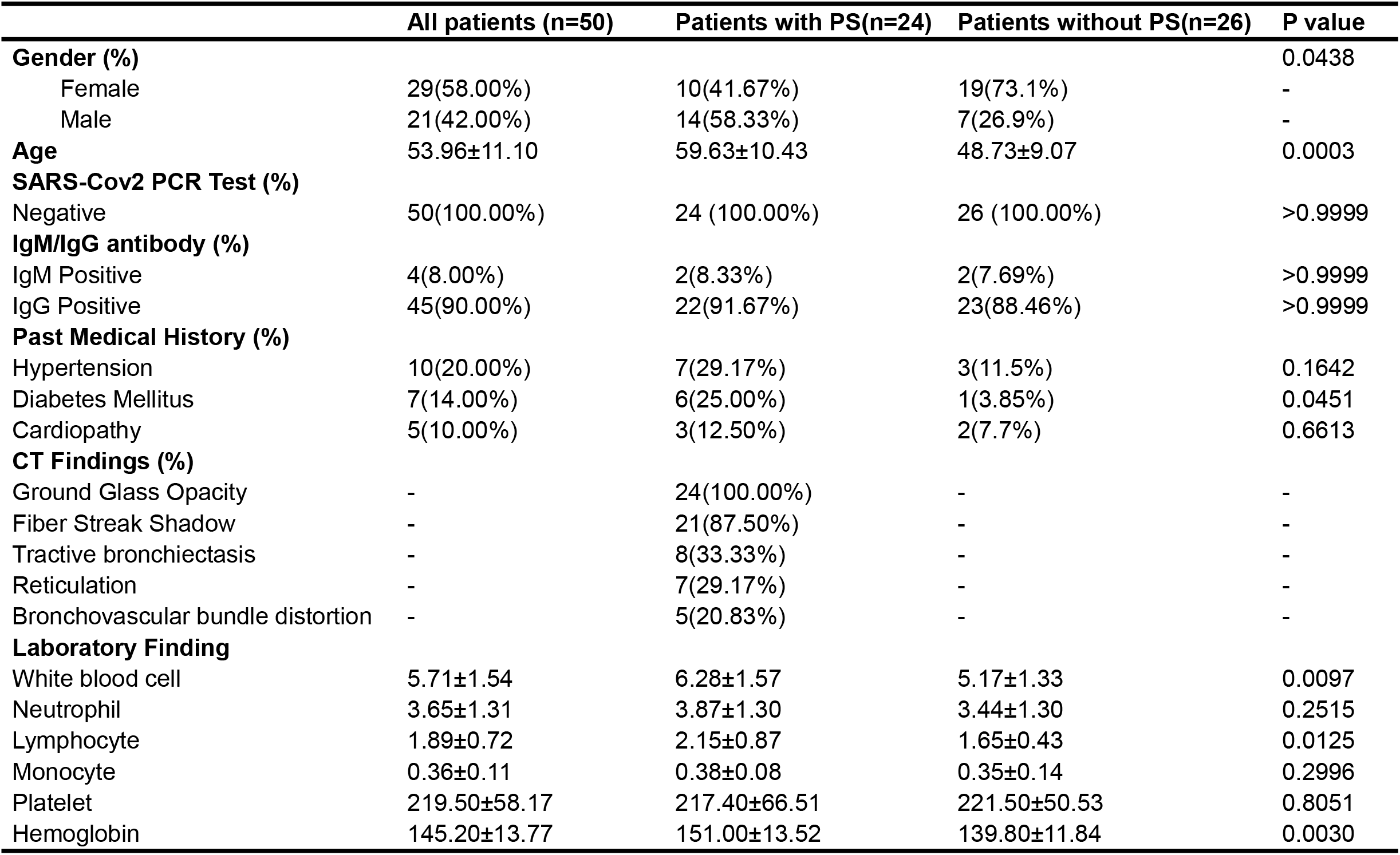
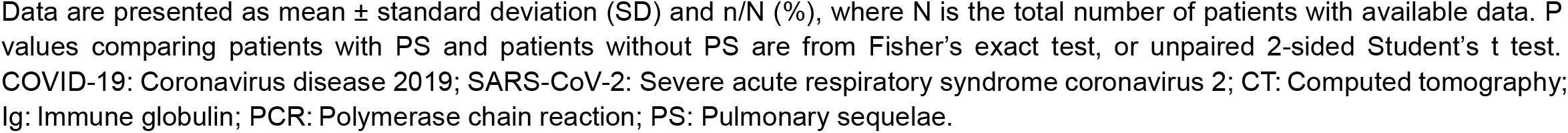
Demographics and clinical characteristics of COVID-19 patients.

|  | **All patients (n=50)** | **Patients with PS(n=24)** | **Patients without PS(n=26)** | **P value** |
| --- | --- | --- | --- | --- |
| **Gender (%)** |  |  |  | 0.0438 |
| Female | 29(58.00%) | 10(41.67%) | 19(73.1%) | - |
| Male | 21(42.00%) | 14(58.33%) | 7(26.9%) | - |
| **Age** | 53.96±11.10 | 59.63±10.43 | 48.73±9.07 | 0.0003 |
| **SARS-Cov2 PCR Test (%)** |  |  |  |  |
| Negative | 50(100.00%) | 24 (100.00%) | 26 (100.00%) | >0.9999 |
| **IgM/IgG antibody (%)** |  |  |  |  |
| IgM Positive | 4(8.00%) | 2(8.33%) | 2(7.69%) | >0.9999 |
| IgG Positive | 45(90.00%) | 22(91.67%) | 23(88.46%) | >0.9999 |
| **Past Medical History (%)** |  |  |  |  |
| Hypertension | 10(20.00%) | 7(29.17%) | 3(11.5%) | 0.1642 |
| Diabetes Mellitus | 7(14.00%) | 6(25.00%) | 1(3.85%) | 0.0451 |
| Cardiopathy | 5(10.00%) | 3(12.50%) | 2(7.7%) | 0.6613 |
| **CT Findings (%)** |  |  |  |  |
| Ground Glass Opacity | - | 24(100.00%) | - | - |
| Fiber Streak Shadow | - | 21(87.50%) | - | - |
| Tractive bronchiectasis | - | 8(33.33%) | - | - |
| Reticulation | - | 7(29.17%) | - | - |
| Bronchovascular bundle distortion | - | 5(20.83%) | - | - |
| **Laboratory Finding** |  |  |  |  |
| White blood cell | 5.71±1.54 | 6.28±1.57 | 5.17±1.33 | 0.0097 |
| Neutrophil | 3.65±1.31 | 3.87±1.30 | 3.44±1.30 | 0.2515 |
| Lymphocyte | 1.89±0.72 | 2.15±0.87 | 1.65±0.43 | 0.0125 |
| Monocyte | 0.36±0.11 | 0.38±0.08 | 0.35±0.14 | 0.2996 |
| Platelet | 219.50±58.17 | 217.40±66.51 | 221.50±50.53 | 0.8051 |
| Hemoglobin | 145.20±13.77 | 151.00±13.52 | 139.80±11.84 | 0.0030 |

Data are presented as mean ± standard deviation (SD) and n/N (%), where N is the total number of patients with available data. P values comparing patients with PS and patients without PS are from Fisher's exact test, or unpaired 2-sided Student's t test. COVID-19: Coronavirus disease 2019; SARS-CoV-2: Severe acute respiratory syndrome coronavirus 2; CT: Computed tomography; Ig: Immune globulin; PCR: Polymerase chain reaction; PS: Pulmonary sequelae.

**FIG 1.**
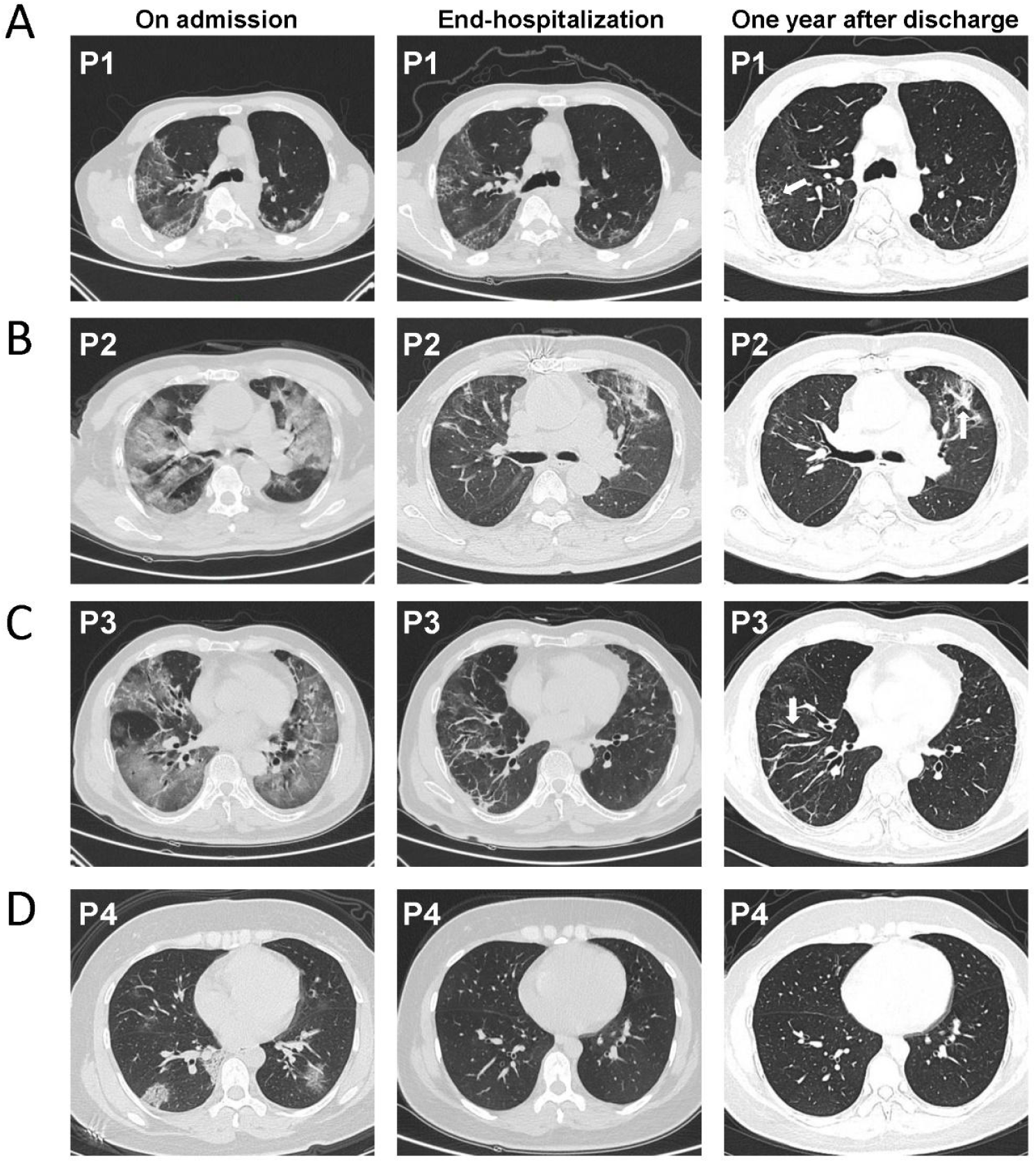
Chest computed tomography scan of four patients across three time periods including on admission, end-hospitalization and one year after discharge. (**A-D**) Chest CT of three time periods showed the change of lung lesions from 3 patients with pulmonary sequelae (P1, P2 and P3) and 1 patient without pulmonary sequelae (P4). (**A**) CT image of a 61-year-old man (P1) showing ground glass opacity (GGO), fiber streak shadow and reticulation one year after discharge. (**B**) CT image of a 77-year-old man (P2) showing GGO, fiber streak shadow and bronchovascular bundle distortion one year after discharge. (**C**) CT image of a 58-year-old man (P3) showing GGO, fiber streak shadow and tractive bronchiectasis one year after discharge. (**D**) CT image of a 36-year-old woman (P4) showing complete resolution of lung lesions one year after discharge.

### NK cells and short-lived effector-like CD8+ T cells accumulate in PPSs

Immunologic disturbance induced by SARS-CoV-2 infection is characterized by lymphopenia in those with acute COVID-19(16). Strikingly, white blood cell and lymphocyte counts were higher in PPSs than NPSs one year after discharge, whereas no significant differences in neutrophil, monocyte and platelet counts were found between the two groups (Table 1). We next analyzed the presence of lymphocyte subsets to obtain an overview of the general distribution in the peripheral blood. The NK cell percentage was significantly higher in PPSs than in NPSs, but there were no significant differences in CD3+ T, CD4+ T, CD8+ T, NKT and γδ T cell percentages (Fig. 2A). A decreased proportion of B cells was detected in PSSs, though the memory B cell frequency was not significantly different (Fig. 2A-B).

**FIG 2.**
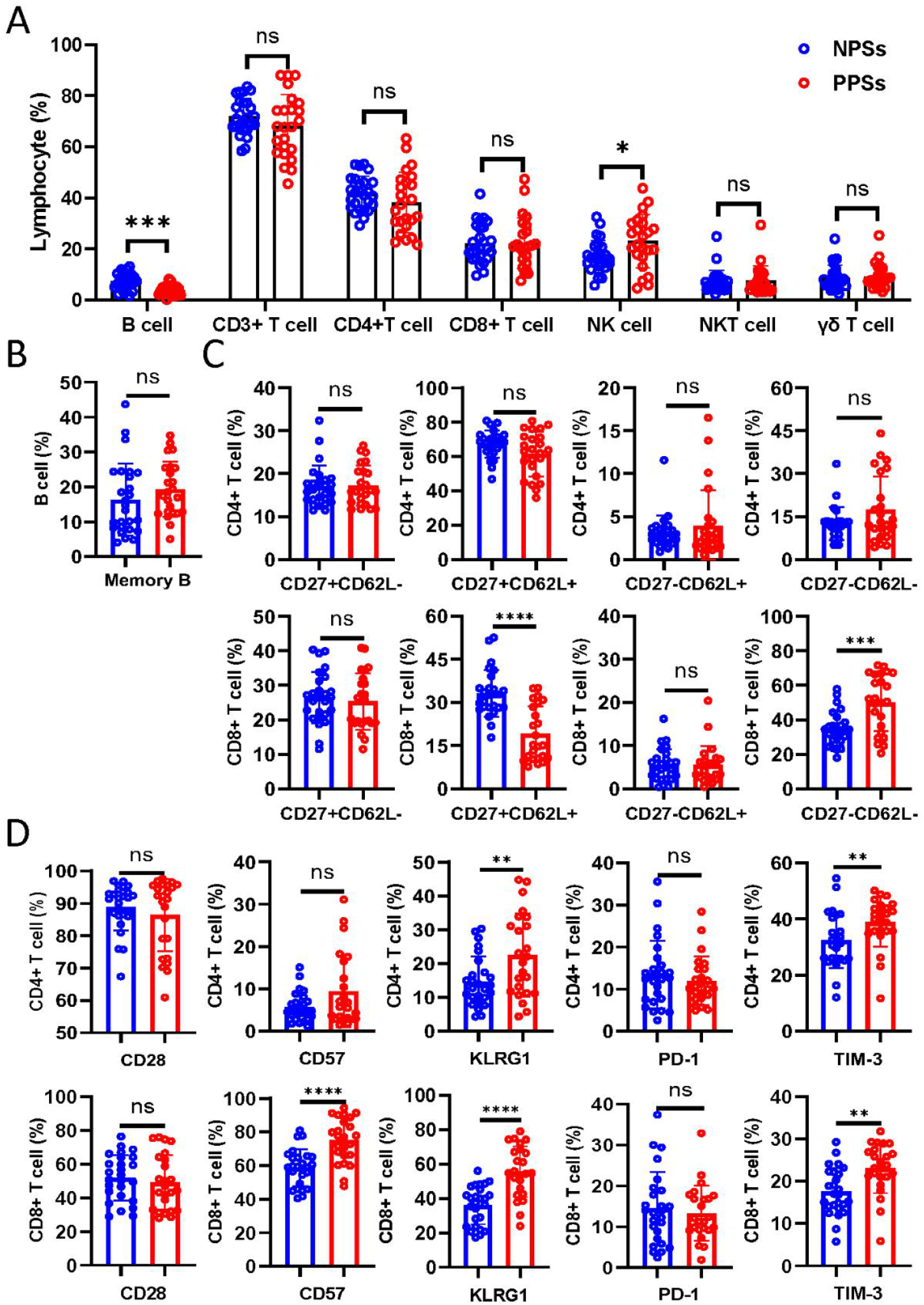
Lymphocyte composition and immunophenotypic characterization of B and T cells in convalescent COVID-19 patients. Circulating lymphocytes from COVID-19 patients with pulmonary sequelae (PPSs, n = 24) and patients without pulmonary sequelae (NPSs, n = 26) were analyzed by multiparameter flow cytometry. (**A**) Relative proportions of CD3+ T, CD4+ T, CD8+ T, NK, NKT, γδ T and B cells between the two groups. (**B**) Relative proportions of memory and naive B cells between the two groups. (**C**) Relative proportions of CD4/CD8+CD27-CD62-, CD4/CD8+CD27+CD62-, CD4/CD8+CD27+CD62+, and CD4/CD8+CD27-CD62+ T cells between the two groups; (**D**) Relative proportions of CD4/CD8+CD28+, CD4/CD8+CD57+, CD4/CD8+KLRG1+, CD4/CD8+PD-1+ T cell and CD4/CD8+ TIM-3+ T cells between the two groups. Data are mean ± SD. The Mann-Whitney U test or unpaired two-tailed Student’s t tests was used to compare the two groups. *p < 0.05, **p < 0.01, ***p < 0.001, ****p < 0.0001.

We next performed immunophenotypic analyses of circulating CD4+ and CD8+ T cells to identify their state of differentiation, exhaustion, and senescence. CD27 and CD62L were used to distinguish maturation and memory subsets in CD4+ and CD8+ T cells. We found no significant differences in CD4+ T cells populations based on CD27 and CD62L expression between PPSs and NPSs (Fig. 2C). CD27-CD62L-T cells represent short-lived effector-like T cells characterized by the enrichment for antigen-experienced and senescent T cells, while CD27+CD62L+ T cells consist of naïve T cells and central memory T cells(17). Importantly, there was a statistically significant increase in the CD8+CD27-CD62L-T cells percentage in PPSs, yet the CD8+CD27+CD62L+ T cell frequency was significantly reduced (Fig. 2C). CD4+ CD57+ T cells percentage tended to be higher in PPSs than in NPSs though there was no statistic difference (Fig. 2D). Higher percentage of CD8+CD57+ T cells was observed in PPSs (Fig. 2D). Moreover, increases of both CD4+ T and CD8+ T cells expressing KLRG-1 and TIM-3 were detected in PPSs, with no changes in the frequencies of CD28-and PD-1-expressing T cells between PPSs and NPSs (Fig. 2D).

### Circulating NK cells and innate-like lymphocytes exhibit distinct phenotypes between PPSs and NPSs

NK, NKT and γδ T cells, large granular lymphocytes with the ability to lyse virally infected cells, are important components of antiviral immunity(18-20). Having established that PPSs displayed an increase in the NK cells percentage, we further analyzed the NK cell phenotype. Importantly, there was a statistically significant increment in NK cells expressing CD57, whereas the frequencies of NKP30- and NKP46-expressing NK cells were significantly reduced in PPSs (Fig. 3A). No changes in the frequencies of CD27-, KLRG1-, PD-1-, TIM-3-, NKB1-, NKG2A- and NKG2D-expressing NK cells were noted between the two groups (Fig. 3A). Focusing on NKT cells, we observed an increase in the fraction of those expressing TIM-3 and NKP30, but the fraction of NKB1+NKT cells was markedly decreased in PPSs (Fig. 3B), suggesting that NKT cells exhibit an activated or exhausted state. However, there were no significant differences in the frequencies of CD27-, CD57-, KLRG1-, PD-1-, NKG2A-, NKG2D- and NKP46-expressing NKT cells between the two groups (Fig. 3B). Moreover, γδ T cells from PPSs overexpressed CD57, KLRG1, TIM-3 and NKP30 (Fig. 3C), which is compatible with a senescent, hyperactivated and exhaustion profile. Downregulation of the inhibitory receptor NKB1 and CD27 in γδ T cells was found in PPSs, but PD-1, NKG2A and NKP46 expressions were unaffected (Fig. 3C).

**FIG 3.**
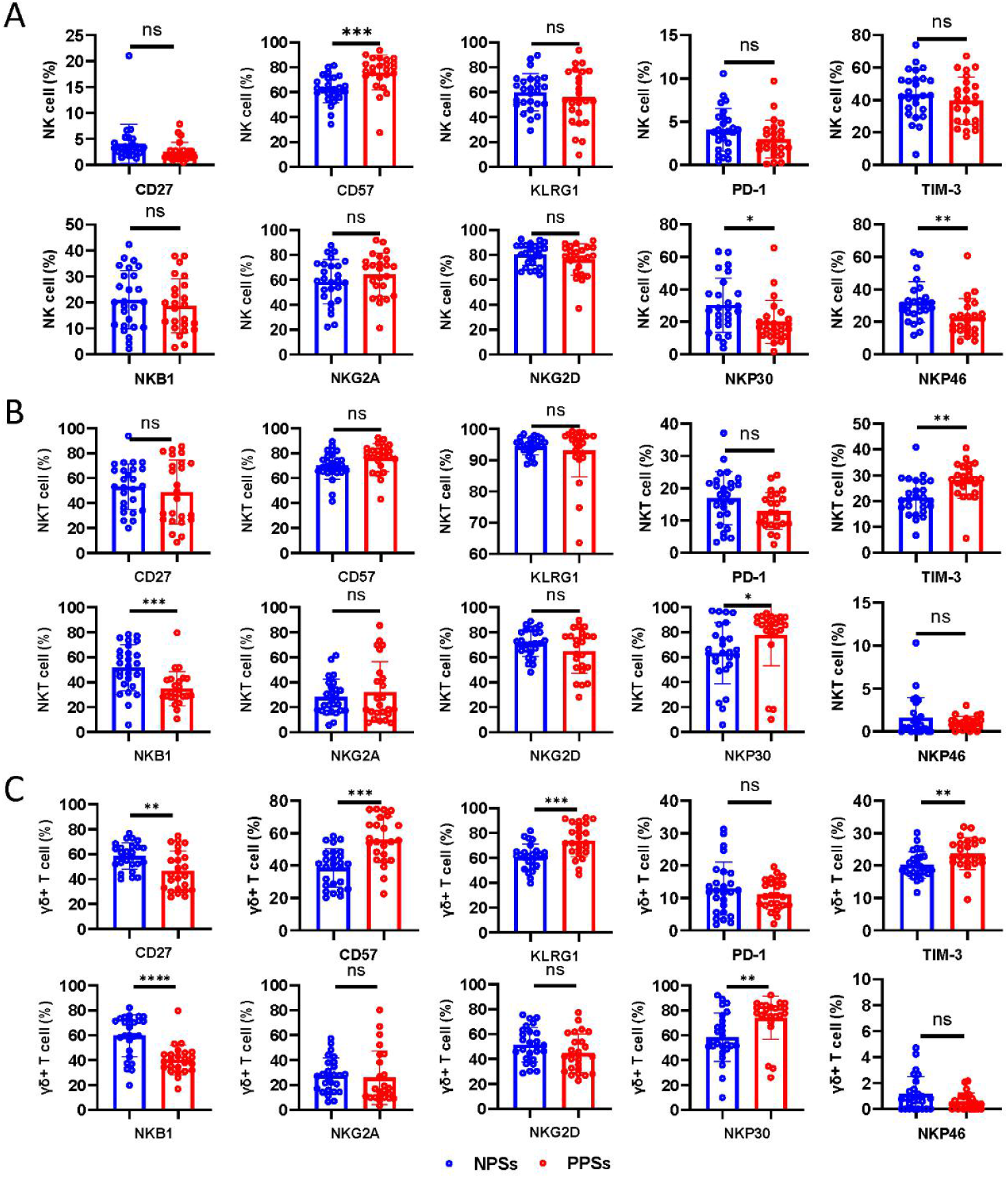
Phenotypical features of NK, NKT and γδT cells in convalescent COVID-19 patients. Circulating NK, NKT and γδT cells from COVID-19 patients with pulmonary sequelae (PPSs, n = 24) and patients without pulmonary sequelae (NPSs, n = 26) were analyzed by multiparameter flow cytometry. (**A**) Relative proportions of CD27, CD57, KLRG1, PD-1, TIM-3, NKB1, NKG2A, NKG2D, NKP30 and NKP46 expression on NK cells between the two groups; (**B**) Relative proportions of CD27, CD57, KLRG1, PD-1, TIM-3, NKB1, NKG2A, NKG2D, NKP30 and NKP46 expression on NKT cells between the two groups; (**C**) Relative proportions of CD27, CD57, KLRG1, PD-1, TIM-3, NKB1, NKG2A, NKG2D, NKP30 and NKP46 expression on γδ T cells between the two groups. Data are mean ± SD. The Mann-Whitney U test or unpaired two-tailed Student’s t tests was used to compare the two groups. *p < 0.05, **p < 0.01, ***p < 0.001, ****p < 0.0001.

### Coexistence of senescence and exhaustion phenotypic signatures in cytokine-secreting cells in PPSs

We next assessed cytokine production in CD3+ T cells after stimulation with PMA and ionomycin and observed that in comparison to NPSs, CD4+ T cells from PPSs overexpressed IL-17A and IFN-γ but not IL-2, IL-4 or TNFα (Fig. 4A). The frequency of IFN-γ expression was higher among CD8+ T cells in PPSs compared with NPSs, with no significant differences in IL-4, IL-17A and TNF-α production in CD8+ T cells between the two groups (Fig. 4B). Nonetheless, CD8+ T cells from PPSs showed downregulated IL-2 expression compared with NPSs (Fig. 4B). We next investigated the degranulation capacity and cytotoxic molecule expression in CD4+ T, CD8+ T, NK, NKT and γδ T cells. Based on functional characterization, CD4+ T, CD8+ T, NK and γδ T cells exhibited upregulation of GZMB and perforin in PPSs compared with NPSs (Fig. 4C-D). However, no significant differences in GZMB and perforin expression in NKT cells were observed (Fig. 4C-D).

**FIG 4.**
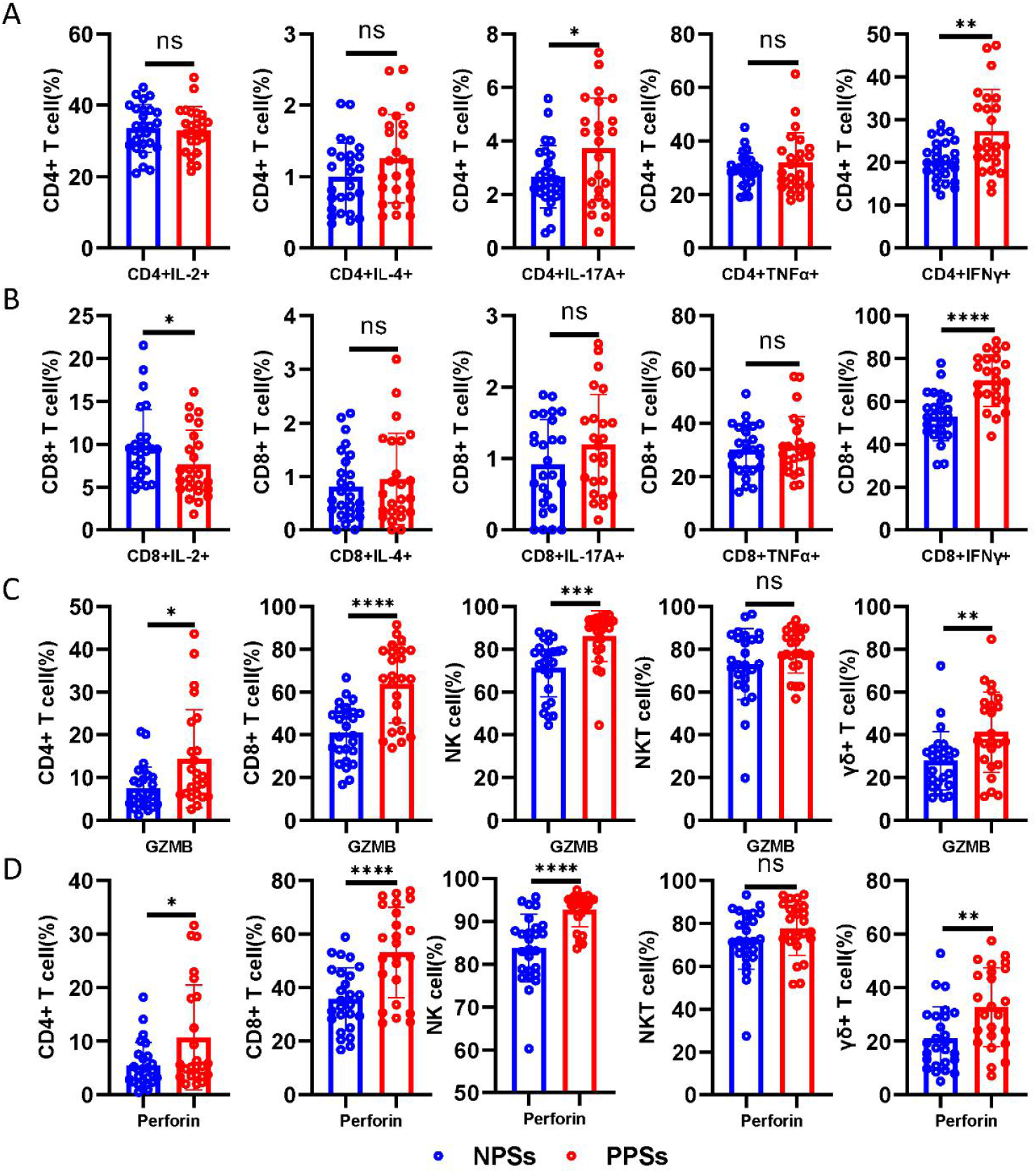
Functional characterization of CD4+ T, CD8+ T, NK, NKT and γδ T cells. Circulating CD4+ T, CD8+ T, NK, NKT and γδ T cells from COVID-19 patients with pulmonary sequelae (PPSs, n = 24) and patients without pulmonary sequelae (NPSs, n = 26) were analyzed by multiparameter flow cytometry. (**A**) Relative proportions of CD4+IL-2+, CD4+IL-4+, CD4+IL-17A+, CD4+IFN-γ+ and CD4+TNF-α+ T cells between the two groups; (**B**) Relative proportions of CD8+IL-2+, CD8+IL-4+, CD8+IL-17A+, CD8+IFN-γ+ and CD8+ TNF-α+ T cells between the two groups. (**C**) Relative proportions of perforin in circulating CD4+ T, CD8+ T, NK, NKT and γδ T cells between the two groups; (**D**) Relative proportions of GZMB in circulating CD4+ T, CD8+ T, NK, NKT and γδ T cells between the two groups. Data are mean ± SD. The Mann-Whitney U test or unpaired two-tailed Student’s t tests was used to compare the two groups. *p < 0.05, **p < 0.01, ***p < 0.001, ****p < 0.0001.

We next used a clustering and visualization strategy to investigate the co-expression of CD27, CD28, PD-1, and CD57 together with cytokines (IL-2, IL-4, IL-17A, TNF-α and IFN-γ) and cytotoxic molecules (GZMB and perforin) (Fig. 5A-B and Supplementary Figure 1H-I). The advantage of this analysis lies in its integration of surface and functional markers at a single-cell level, providing an improved understanding of their high-dimensional relationship. CD4+IL-2+ T, CD4+IL-4+ T and CD4+IL-17A+ T cells hardly expressed CD57 (Fig. 5A). Phenotypic characterization further demonstrated that CD4+ T cells expressing IL-2, IL-4 and IL-17A were prominent among the CD28+ T cells (Fig. 5A). According to representative scatter plot, IFN-γ could be secreted by CD4+CD57+ T cells and CD4+CD57-T cells; furthermore, CD4+CD57+IFN-γ+ T cells were enriched among CD4+CD28-T cells (Fig. 5A). As observed in CD4+ T cells, the co-expression of IL-2, IL-4, and IL-17A together with CD28 and CD57 could be similarly observed in CD8+ T cells (Supplementary Fig. 1H). Next, we investigated whether PD-1 expression correlates with cytokine production in the T cell population. We found that IL-2, IL-4, IL-17A, TNF-α and IFN-γ expressions were mainly distributed in PD-1-T cells (Fig. 5A and Supplementary Fig. 1H). We summarized these data in a heatmap and exhibited the distributions of IL-2, IL-4, IL-17A, TNF-α and IFN-γ expression together with CD28, CD57 and PD-1 between the two groups (Fig. 5C).

**FIG 5.**
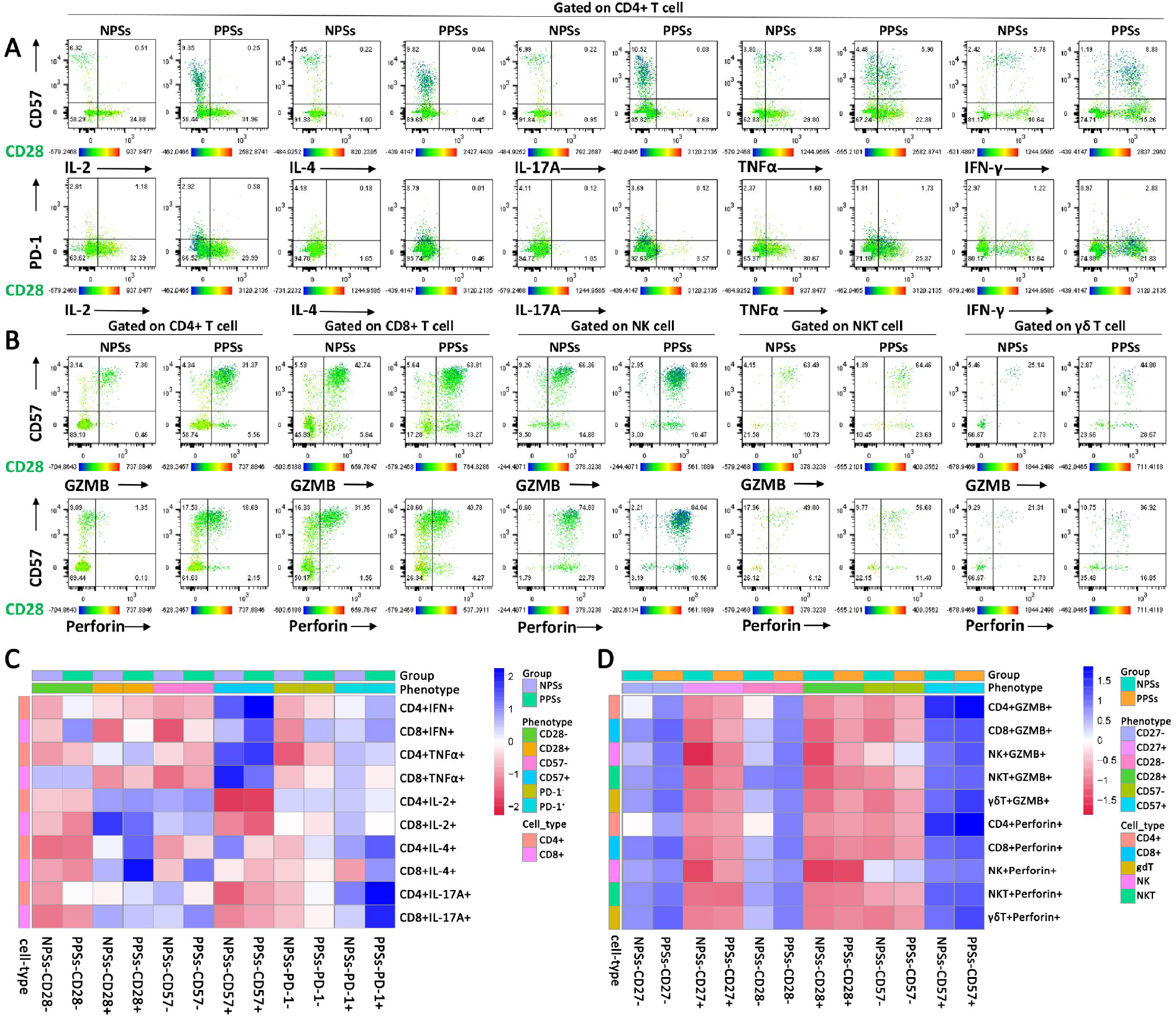
Senescence and exhaustion phenotypes coexist in cytokine-secreting CD4+ T, CD8+ T, NK, NKT and γδ T cells. The co-expressions of CD27, CD28 and PD-1 together with cytokines (IL-2, IL-4, IL-17A, TNF-α and IFN-γ) in CD4+ T and CD8+ T cells from COVID-19 patients with pulmonary sequelae (PPSs, n = 24) and patients without pulmonary sequelae (NPSs, n = 26) were analyzed by multiparameter flow cytometry. (**A**) A bidimensional map obtained from flow cytometric data displaying the co-expression of CD27, CD28 and PD-1 together with cytokines (IL-2, IL-4, IL-17A, TNF-α and IFN-γ) in CD4+ T cell. Supplementary Figure 1H displayed the co-expression of CD27, CD28 and PD-1 together with cytokines (IL-2, IL-4, IL-17A, TNF-α and IFN-γ) in CD8+ T cell. The co-expressions of CD27, CD28 and 57 together with granzyme B (GZMB) and perforin in circulating CD4+ T, CD8+ T, NK, NKT and γδ T cells from COVID-19 patients with pulmonary sequelae (PPSs, n = 24) and patients without pulmonary sequelae (NPSs, n = 26) were analyzed by multiparameter flow cytometry. (**B**) A bidimensional map obtained from flow cytometric data displaying the co-expression of CD28 and CD57 together with GZMB and perforin in CD4+ T, CD8+ T, NK, NKT and γδ T cells. Supplementary Figure 1I displayed the co-expression of CD27 and CD57 together with GZMB and perforin in CD4+ T, CD8+ T, NK, NKT and γδ T cells. (**C**) Heatmap represents the percentage of cells in CD27+, CD27-, CD28+, CD28-, PD-1+ and PD-1-clusters that express IL-2, IL-4, IL-17A, TNF-α and IFN-γ in CD4+ T and CD8+ T cells for PPSs and NPSs. (**D**) Heatmap represents the percentage of cells in CD27+, CD27-, CD28+, CD28-, CD57+ and CD57-clusters that express GZMB and perforin in CD4+ T, CD8+ T, NK, NKT and γδ T cells for PSSs and NPSs.

We next investigated the expression of CD27, CD28, and CD57 together with cytotoxic molecules in CD4+ T, CD8+ T, NK, NKT and γδ T cells (Fig. 5B). According to representative scatter plot, GZMB and perforin expression were higher among CD4+CD57+ T cells than CD4+CD57-T cells in both groups; furthermore, CD4+CD57+GZMB+ T cells and CD4+CD57+perforin+ T cells were mainly enriched among CD27-cells and CD28-cells (Fig. 5B and Supplementary Fig. 1I). As observed in CD4+ T cells, GZMB and perforin expression in CD8+ T, NK, NKT and γδ T cells were similarly enriched among CD57+ cells, CD28-cells and CD27-cells (Fig. 5B and Supplementary Fig. 1I). We summarized the data in a heatmap and exhibited the distributions of GZMB and perforin expressions together with CD27, CD28, and CD57 between the two groups (Fig. 5D).

### Comparable immunosuppressive cells between PPSs and NPSs

Immunosuppressive cells served as the main mechanisms for maintaining immune homeostasis. However, there is considerable controversy regarding whether immunosuppressive cells promote or constrain the formation of fibrosis induced by pathogenic T cells(21-24). We next explored the distribution of immunosuppressive cells in recovered patients. After lysis of the erythrocytes, we directly used cell surface markers to stain monocytic myeloid-derived suppressor cells (M-MDSCs, HLA-DR-/lowCD33+CD11b+CD14+), granulocytic MDSCs (G-MDSCs, HLA-DR-/low CD33+CD11b+CD14-CD15+), regulatory T cells (Tregs, CD4+CD127-/lowCD25+) and regulatory B cells (Bregs, CD19+CD24+CD38+)(25, 26). Certainly, it is not stringent to define G-MDSCs by using this method. Patients with acute COVID-19 produce emergency myelopoiesis-generating immunosuppressive myeloid cells(27). But, M-MDSCs and G-MDSCs frequencies did not differ between PPSs and NPSs (Fig. 6A-B). A decrease in HLA-DR on monocytes in acute COVID-19 is associated with severe respiratory failure(28). We did not observe any significant difference in HLA-DR expression on monocyte between the two groups one year after discharge (Data not shown). Additionally, Tregs and Bregs showed no significant differences among the groups (Fig. 6C-D). Together, our data indicate that immunosuppressive cells may be insufficient to constrain the robust activation of a variety of immune cells in PPSs.

**FIG 6.**
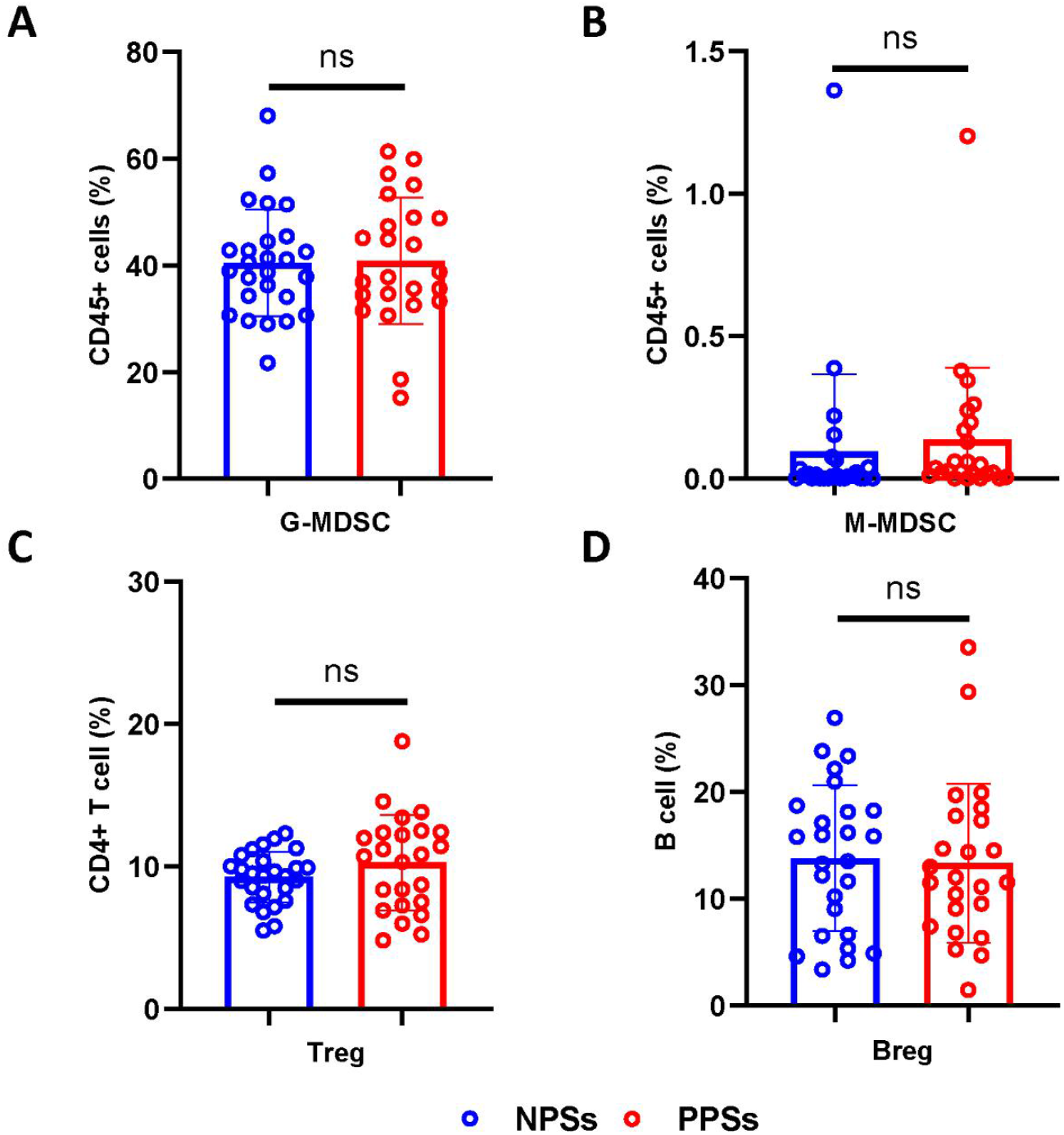
Distribution of immunosuppressive cells between patients with PS and patients without PS. Circulating G-MDSCs, M-MDSCs, Tregs, and Bregs from COVID-19 patients with pulmonary sequelae (PPSs, n = 24) and patients without pulmonary sequelae (NPSs, n = 26) were analyzed by multiparameter flow cytometry. (**A**) Relative proportions of G-MDSCs between the two groups; (**B**) Relative proportions of M-MDSCs between the two groups. (**C**) Relative proportions of Tregs frequencies between the two groups; (**D**) Relative proportions of Bregs between the two groups. Data are mean ± SD. The Mann-Whitney U test or unpaired two-tailed Student’s t tests was used to compare the two groups. *p < 0.05, **p < 0.01, ***p < 0.001, ****p < 0.0001.

### The interrelation of immune cells and its association with clinical features in COVID-19

Having established that senescent signatures coexist within cytotoxic molecule-secreting cells, we further investigated the interrelation of immune cells in convalescent patients with COVID-19 (Fig. 7A). Correlation analysis verified that T cells secreting cytotoxic molecules correlated negatively with CD27+CD62L+ T cells and CD28+ T cells but positively with short-lived effector-like CD27-CD62L-T cells, CD57+ T cells and KLRG1+ T cells (Fig. 7A). CD57 and KLRG-1 are thought to be associated with T cell senescence and also serves as a marker of cytotoxic function(29). Importantly, IL-2 expression within CD8+ T cells correlated negatively with CD8+CD27-CD62L-T cells, CD8+CD57+ T cells, CD8+perforin+ T cells and CD8+GZBM+ T cells (Fig. 7A). The downregulated IL-2 expression in CD8+ T cells from PPSs further verifies that these patients retained large amounts of short-lived like CD8+ T cells that abundantly secrete cytotoxic molecules. Moreover, the expression of perforin and GZMB within NK cells were also positively associated with effector molecule CD57 but inversely with PD-1, NKP30 and NKP46 expression (Fig. 7A). Additionally, GZMB and perforin expression exhibited intrinsic positive correlations among CD4+ T, CD8+ T, NK, NKT and γδ T cells, suggesting that SARS-CoV-2 infection may simultaneously trigger robust activation of a variety of immune cells. More detailed information is displayed in the correlation heatmap depicted in (Fig. 7A).

**FIG 7.**
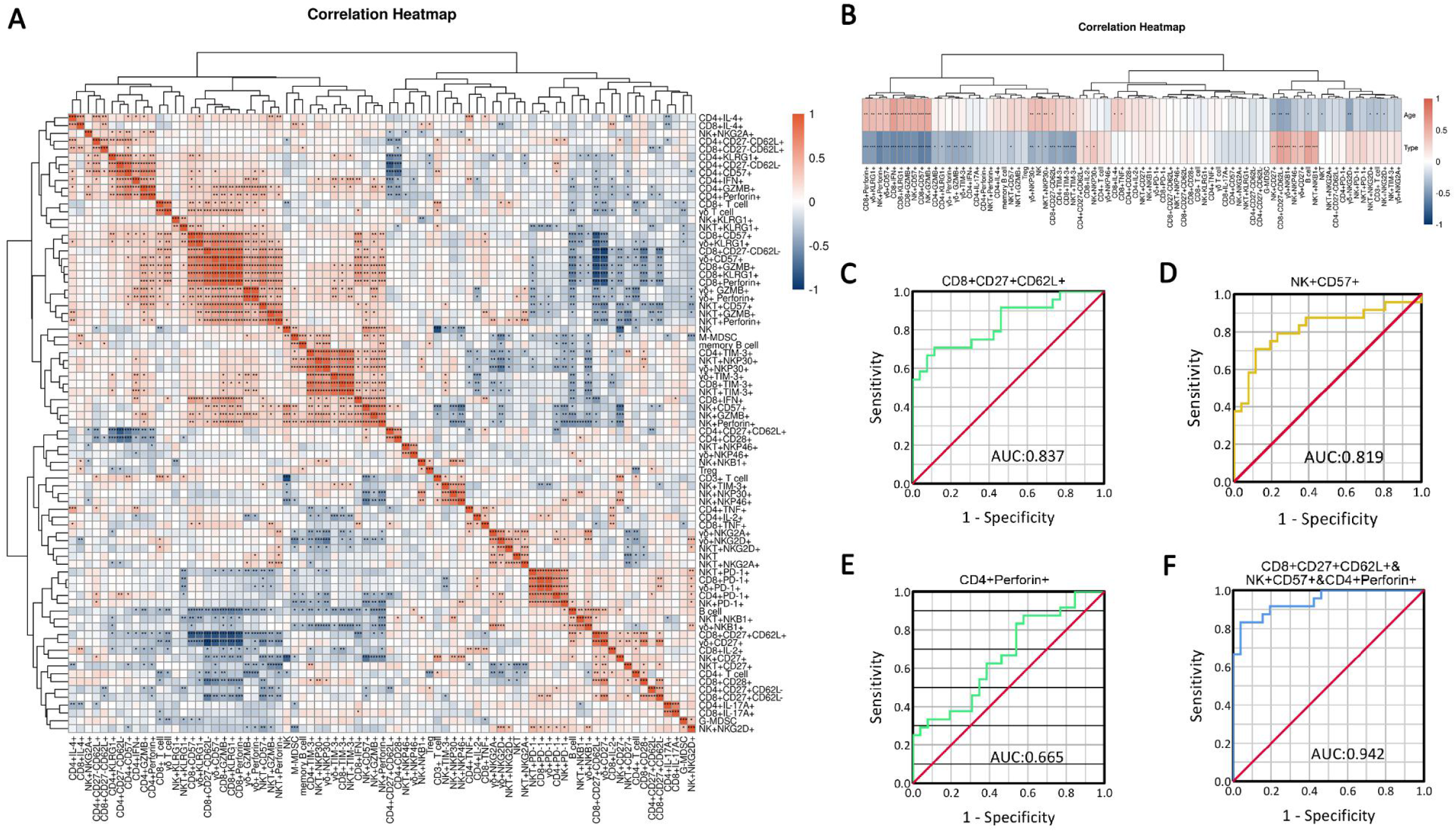
The interrelation of immune cells and its correlation with clinical features. (**A**) Correlation heatmap exhibited the interrelation of immune cells in all recruited patients with COVID-19. (**B**) Correlation heatmap exhibited the correlation of immunological parameters with age and disease type (two types: patients with pulmonary sequelae and patients without pulmonary sequelae). Spearman’s rank coefficient was used to determine correlations between two variables. The three most contributing variables (CD8+CD27+CD62+ T cells, CD57+NK cells and CD4+perforin+ T cells) were identified by multivariate logistic regression analyses. (**C-F**) ROC curves were calculated for these selected parameters by using SPSS.

We next sought to determine whether the phenotypic and functional features of immune cells correlate with age and disease type in COVID-19. Correlation analysis suggested that age significantly influenced the phenotypic and functional features of immune cells (Fig. 7B). Senescent and short-lived like CD8+ T cells, CD8+ T cells secreting perforin/granzyme-B/IFN-γ, NK cells secreting perforin/granzyme-B, NKP30+NKT, TIM-3+NKT, NKP30+γδ T, KLRG1+γδ T and CD57+γδ T cells percentages correlated positively with age, whereas B cells, CD3+ T cells, NKT cells, CD8+CD27+CD62L+ T cells, NKB1+γδ T cells, NKG2D+γδ T cells, CD27+NK cells, NKG2D+NK cells and NKG2D+NKT cells frequencies correlated negatively with age (Fig. 7B). Furthermore, correlation analysis of immunological parameters in PPSs and NPSs revealed that senescent, exhausted, GZMB and perforin secreting immune cells correlated positively with PS (Fig. 7B). Additionally, CD8+CD27-CD62L-T cells, NK cells, M-MDSCs, NKP30+γδ T cells, NKP30+NKT cells, CD4+ IFN-γ+ and CD8+IFN-γ+ T cells percentages correlated positively with PS (Fig. 7B).

Next, we examined the possibility of using the above-mentioned parameters as prognostic factors for identifying determinants for residual lesions in COVID-19 patients. Multivariate logistic regression analyses identified CD8+CD27+CD62L+ T cells (odds ratio [OR]: 0.738; 95% CI: 0.590-0.924; p = 0.008), CD57+NK cells (OR: 1.181; 95% CI: 1.038-1.343; p = 0.012), and CD4+perforin+ T cells (OR:1.153; 95% CI: 0.953-1.396; p = 0.143) as independent predictors for residual lung lesions (Table 2). Additionally, receiver operating characteristic curve were carried out to assess the capacity of the three cell populations to differentiate disease type (PPSs and NPSs). The cutoff values (sensitivity and specificity) are as follows: CD8+CD27+ CD62L+ T cells: 26.045% (0.885% and 0.708%), CD57+NK cells: 74.095% (0.708% and 0.885%), CD4+perforin+ T cells: 3.245% (0.875% and 0.423%). The AUC value of the combination of the three cells (CD57+NK cells, CD4+perforin+ T cells and CD8+CD27+CD62L+ T cells) was highest (0.942) (Fig. 7C-F).

**Table 2.** Univariate and multivariate logistic regression analyses of PS.

|  | Univariate logistic regression |  |  |  | Multivariate logistic regression |  |  |  |
| --- | --- | --- | --- | --- | --- | --- | --- | --- |
|  | OR | 95% CI |  | P value | OR | 95% CI |  | P value |
|  |  | Low | Up |  |  | Low | Up |  |
| Age (year) | 1.117 | 1.043 | 1.197 | 0.002 |  |  |  |  |
| B cell (%) | 0.590 | 0.435 | 0.802 | 0.001 |  |  |  |  |
| CD8+CD27+CD62L+ (%) | 0.834 | 0.754 | 0.922 | <0.001 | 0.738 | 0.590 | 0.924 | 0.008 |
| CD8+CD27-CD62L- (%) | 1.083 | 1.031 | 1.138 | 0.001 |  |  |  |  |
| NK (%) | 1.090 | 1.012 | 1.174 | 0.023 |  |  |  |  |
| CD4+KLRG1+ (%) | 1.082 | 1.016 | 1.153 | 0.015 |  |  |  |  |
| CD4+TIM-3+ (%) | 1.082 | 1.010 | 1.158 | 0.024 |  |  |  |  |
| CD8+CD57+ (%) | 1.114 | 1.047 | 1.185 | 0.001 |  |  |  |  |
| CD8+KLRG1+ (%) | 1.122 | 1.051 | 1.198 | 0.001 |  |  |  |  |
| CD8+TIM-3+ (%) | 1.186 | 1.055 | 1.332 | 0.004 |  |  |  |  |
| NK+CD57+ (%) | 1.096 | 1.033 | 1.164 | 0.003 | 1.181 | 1.038 | 1.343 | 0.012 |
| NKT+TIM-3+ (%) | 1.155 | 1.043 | 1.279 | 0.005 |  |  |  |  |
| CD57+γδT (%) | 1.105 | 1.043 | 1.171 | 0.001 |  |  |  |  |
| KLRG1+γδT (%) | 1.098 | 1.034 | 1.165 | 0.002 |  |  |  |  |
| TIM-3+γδT (%) | 1.196 | 1.036 | 1.381 | 0.014 |  |  |  |  |
| NK+NKP30+ (%) | 0.953 | 0.912 | 0.996 | 0.031 |  |  |  |  |
| NK+NKP46+ (%) | 0.930 | 0.877 | 0.987 | 0.017 |  |  |  |  |
| NKT+NKB1+ (%) | 0.939 | 0.900 | 0.979 | 0.003 |  |  |  |  |
| NKB1+γδT (%) | 0.920 | 0.878 | 0.965 | 0.001 |  |  |  |  |
| NKP30+γδT (%) | 1.049 | 1.011 | 1.087 | 0.010 |  |  |  |  |
| CD4+IFN+ (%) | 1.151 | 1.038 | 1.276 | 0.008 |  |  |  |  |
| CD4+IL-17A+ (%) | 1.550 | 1.046 | 2.296 | 0.029 |  |  |  |  |
| CD8+IFN+ (%) | 1.133 | 1.057 | 1.215 | <0.001 |  |  |  |  |
| CD4+GZMB+ (%) | 1.118 | 1.016 | 1.230 | 0.022 |  |  |  |  |
| CD4+Perforin+ (%) | 1.112 | 1.009 | 1.226 | 0.033 | 1.153 | 0.953 | 1.396 | 0.143 |
| CD8+GZMB+ (%) | 1.089 | 1.038 | 1.142 | <0.001 |  |  |  |  |
| CD8+Perforin+ (%) | 1.083 | 1.033 | 1.135 | 0.001 |  |  |  |  |
| NK+GZMB+ (%) | 1.113 | 1.040 | 1.192 | 0.002 |  |  |  |  |
| NK+Perforin+ (%) | 1.304 | 1.128 | 1.507 | <0.001 |  |  |  |  |
| GZMB+γδT (%) | 1.054 | 1.012 | 1.097 | 0.012 |  |  |  |  |
| Perforin+γδT (%) | 1.067 | 1.018 | 1.118 | 0.007 |  |  |  |  |
| CD27+γδT (%) | 0.938 | 0.896 | 0.982 | 0.006 |  |  |  |  |

## DISCUSSION

Although a tremendous global effort by the scientific community has greatly improved our understanding of COVID-19, the immunopathogenesis of PS remains to be elucidated. Here, we describe the circulating immune landscape of COVID-19 patients with PS compared with those without PS. Residual lesions in PPSs were mainly GGO and fiber streak shadow. Our study demonstrates that there existed significantly divergent in immunological characteristics between PPSs and NPSs one year after discharge. Immunological signatures in PPSs were predominated by the accumulation of CD4+ T cells secreting IL-17A and short-lived effector-like CD8+ T cells with excessive secretion of IFN-γ/granzyme-B/perforin. NK cells were characterized by excessive secretion of granzyme-B and perforin and downregulation of NKP30 and NKP46; NKT and γδ T cells demonstrated highly activated and exhausted states in PPSs. Overall, we observed robust activation of a variety of immune cells in response to SARS-CoV-2 infection and specific features unique to COVID-19 with PS and hyperinflammation, providing a base for understanding the role of cellular immunity in patients with COVID-19.

Over 80% of patients with COVID-19 experience lymphopenia and exhibit drastically reduced numbers of various lymphocyte subsets, including CD4+ T, CD8+ T, B, γδ T and NK cells, especially in the peripheral blood of those with severe COVID-19 during the acute phase(16, 30-34). Early-stage lymphopenia is thought to correlate with lymphocyte chemotaxis in COVID-19(35, 36). To our surprise, lymphocyte counts were higher in PPSs than in NPSs one year after discharge, suggesting that lymphocytes may exit inflamed tissues and undergo clonal expansion after the resolution of SARS-CoV-2 infection. In general, clonal expansion of both innate and adaptive lymphocytes is a critical process for host defense via amplification of lymphocytes specific to the invading pathogen. The NK cell percentage was significantly higher in PPSs than in NPSs, whereas the B cell frequency was lower in the former, suggesting that NK cells underwent more significant clonal expansion. A previous study found that decreased B cells were associated with prolonged viral RNA shedding from the respiratory tract in COVID-19(37). Nevertheless, the reason for the obvious expansion of NK cells following SARS-CoV-2 clearance in PPSs remained to be elucidated. Prolonged viral RNA shedding may induce clonal expansion of NK cells in an antigen-specific manner, similar to the response to cytomegalovirus infection to acquiring memory features(38). Since reverse-transcribed SARS-CoV-2 RNA can integrate into the genome of cultured human cells and can be expressed in patient-derived tissues(39), there might also be latent SARSCoV-2 virus in the reservoir cells in PPSs that could lead to the prolonged expansion of NK cells.

Excessive activation of proinflammatory immune cells can lead to enhanced inflammation and injury during pulmonary viral infection(40). COVID-19 promotes cell polarization of naive and memory cells to effector, cytotoxic, exhausted and regulatory cells, along with increased late NK cells, and induces gene expression related to inflammation and cellular senescence(41). By examining the phenotypic characteristics of T cells, we observed a low percentage of CD8+CD27+CD62L+ T cells but high level of CD8+CD27-CD62L-T cells in PSSs. T cells in PSSs displayed an overall exhausted and senescent phenotype, with overexpression of CD57, KLRG-1 and TIM-3. Functional analysis further revealed that upregulation of degranulation capacity and cytotoxic molecules in CD4+ and CD8+ T cells in PPSs compared with NPSs. Our findings are in line with published papers(42-44). SARS-CoV-2 infection may induce a cytotoxic response, characterized by simultaneous production of GZMB and perforin in T cells, NK cells and γδ T cells in PPSs. Excessive activation of cytotoxic T, NK and γδ T cells is not protective but rather drives pulmonary damage after SARS-CoV-2 infection.

Emerging evidence suggests that COVID-19 survivors have impaired lung function, with the development of pulmonary fibrosis(2, 14). Tissue-resident CD8 + T cells drive age-associated chronic lung sequelae after viral pneumonia(45). Besides, clonally expanded tissue-resident memory-like Th17 cells may interact with pro-fibrotic macrophages and cytotoxic CD8+ T cells leading to the formation of pulmonary fibrosis(4). We observed that peripheral blood CD4+ T cells overexpressed IL-17A/IFN-γ and that CD8+ T cells overexpressed IFN-γ/GZMB/ perforin in PSSs. CD103high Tregs can constrain lung fibrosis induced by CD103low tissue-resident pathogenic CD4+ T cells with higher production of effector cytokines, such as IL-4, IL-5, IL-13, IL-17A and IFN-γ(23). However, immunosuppressive cells were comparable between PPSs and NPSs in our study, indicating that these cells may be insufficient to constrain the robust activation of a variety of immune cells in PSSs. Considering that Tregs are equally important to prevent inflammation-induced tissue damage during acute infections and to promote tissue repair, the scholars suggest that Tregs-based strategies could be considered for COVID-19 patient management(46). Hence, immune intervention may be one of the effective treatment measures to reduce the occurrence of PS after the resolution of SARS-CoV-2 infection.

Several other important findings emerged from our data. First, IL-2+ T cells, IL-4+ T cells and IL-17A+ T cells hardly expressed CD57 but were enriched among CD28+ cells, indicating that autocrine cytokines may provide a tonic signal that inhibits senescence. Furthermore, CD4+ T cells, CD8+ T cells, NK cells, NKT cells and γδ T cells secreting GZMB and perforin were enriched among CD57+ cells, CD28-cells and CD27-cells. Although peripheral blood CD57+ T cells exhibit phenotypic and functional features of terminally differentiated effector cells, these cells display enhanced cytotoxic function(47-49). Moreover, GZMB and perforin expression exhibited intrinsic positive correlations among CD4+ T cells, CD8+ T cells, NK cells, NKT cells and γδ T cells, suggesting that SARS-CoV-2 infection may simultaneously activate a variety of immune cells. In addition, we also noted downregulated expression of NKP30 and NKP46 in NK cells, despite excessive secretion of perforin and GZMB in PPSs. Combined with previous research showing reduced surface expression of NKP30 and NKP46 on adaptive memory NK cells(50), downregulated expression of NKP30 and NKP46 may act as a protective mechanism against tissue damage induced by excessive secretion of perforin and GZMB. Certainly, SARS-CoV-2 may escape the killing of NK cells and damage lung tissue due to downregulated expression of NKP30 and NKP46.

Although our study confirms some findings and provides new data on the innate and adaptive immune landscape of patients with PS who have recovered from COVID-19 one year after discharge, we recognize limitations that might be overcome with larger sample sizes and matched control populations. Furthermore, there is a lack of understanding of the phenotype and function of immune cells from the lungs, which may directly participate in the formation of PS. Hence, the hierarchy of immunodominant circulating blood immune cells may not exactly reflect immunophenotypic features in the lungs. In summary, our study first shows significant differences in immunological characteristics between PPSs and NPSs one year after discharge. Although the detailed mechanisms by which cellular immunity participates in the development of PS remain to be investigated, our in-depth analysis of immunological profiling contributes to our understanding of the immunopathogenesis of COVID-19, facilitating the tailoring of more effective and proactive therapies for these patients.

## MATERIALS AND METHODS

### Study design and participants

In order to determine the immunopathogenesis of residual lung lesions in COVID-19 survivors one year after discharge, a total of 50 convalescent patients were recruited at union hospitals. At the visit, routine blood test, chest computed tomography (CT) scans, and nucleic acid test and antibody detection for SARS-CoV-2 were performed for each participant. The patients were classified as 24 PPSs and 26 NPSs according to radiological characteristics. Patients who reached complete radiological resolution were regarded as NPSs. Complete radiological resolution was defined as the absence of any chest radiographic abnormality potentially related to infection(51). PS including residual GGO, fibrous stripe shadow, tractive bronchiectasis, reticulation and bronchovascular bundle distortion were evaluated by two radiologists. During our recruitment process, we excluded participants with the underlying chronic lung diseases and cancers. This study was conducted in accordance with the Declaration of Helsinki and was approved by the Ethics Committee of Union Hospital, Tongji Medical College, Huazhong University of Science and Technology (#2020/0004), and written informed consents were obtained from all participants.

### Detection of SARS-CoV-2 mRNA and serum SARS-CoV-2 IgG and IgM

SARS-CoV-2 RNA was detected by reverse transcription-polymerase chain reaction (RT-PCR). Total nucleic acid extraction from nasopharyngeal specimens was performed using the QIAamp RNA Viral Kit (Qiagen), and two sets of primers were taken for two target genes (ie, open reading frame 1ab [ORF1ab] and nucleocapsid protein [N]). Detection of serum SARS-CoV-2 IgG and IgM antibodies was evaluated by IgM/IgG antibody detection kit (Abbott Laboratories, Inc).

### Multiparametric flow cytometric analysis

Peripheral blood (200µL) from convalescent patients with COVID-19 was added into a tube and 2 mL of red-cell lysis buffer was added and incubated for 10 minutes. After lysis, the sample was washed twice with PBS containing 1% FBS. Immune cells were surface-stained with fluorochrome-conjugated antibodies. The samples were incubated with antibodies for 15 minutes at 4 °C. Cells were resuspended in PBS and washed at 400 *g* for 6 minutes. The specimens were immediately valuated by flow cytometry.

For cytotoxic molecule detection, we isolated peripheral blood mononuclear cells (PBMCs) from heparinized blood by Ficoll-Hypaque gradient centrifugation (Pharmacia, Uppsala, Sweden). For surface staining, PBMCs were washed twice with PBS containing 1% FBS and stained with fluorochrome-conjugated antibodies. Intracellular staining for granzyme B (GZMB) and perforin was performed after cell fixation and permeabilization (eBioscience), and then intracellular proteins were labeled with the corresponding antibodies conjugated with fluorescent molecules according to the manufacturer’s instructions.

A list of the antibodies used is provided in Supplementary Table 1, and the gating strategy is presented in Supplementary Fig. 1. Flow cytometry was performed using a BD LSRFortessa X-20 (BD Biosciences), and data were analyzed with FlowJo V10 software.

### Cytokine production assays

PBMCs were cultured in RPMI1640 supplemented with 10% FBS, and cytokine production assays were performed after lymphocytes were stimulated with polymethyl acrylate (PMA, 50 ng/mL) and ionomycin (1 μM) in the presence of Golgi-Stop. After 5 hours at 37 °C, the cells were stained with fluorochrome associated antibodies specific for surface molecules; next, the cells underwent fixation and permeabilization for intracellular staining with antibodies specific for the following intracellular proteins: IL-2, IL-4, IL-17A, IFN-γ and TNF-α.

### Statistical analysis

The Shapiro-Wilk test was used to evaluate the distribution of variance. Continuous variables with normally and nonnormally distributed data were assessed using unpaired two-tailed Student’s t tests or Mann-Whitney U test, respectively. The Fisher’s exact test was applied to examine categorical variables. Spearman’s rank coefficient was used to determine correlations between two variables. Multivariate logistic regression analyses were performed to identify the independent predictive factors of residual lesions. The final model was determined using stepwise logistic regression, with significance level for selection set at p = 0.05. The optimum cut-off values were defined based on their maximum Youden index (sensitivity+ specificity-1). All tests were 2-sided, and significance levels were set to p < 0.05 (*), p < 0.01 (**), p < 0.001 (***), p < 0.0001 (****) and ns means not significant. All statistical data were analyzed using SPSS version 25.0 Statistical Software (Chicago, IL, USA), GraphPad Prism 8 software (GraphPad Software, La Jolla, California) or R software Version 4.0.2 (Institute for Statistics and Mathematics, Vienna, Austria).

## ACKNOWLEDGMENTS

We would like to thank all participating patients for their support and cooperation as well as our colleagues from Imaging Department of Union hospital. This work was supported by funds from the Key Special Project of Ministry of Science and Technology, China (No.2020YFC0845700), Scientific Research Projects of Chinese Academy of Engineering (NO.2020-KYGG-01-07) and Fundamental Research Funds for the Central Universities (NO.2020kfyXGYJ029).

